# Suppression of hepatic amino acid catabolism by the human-specific lncRNA hLMR1

**DOI:** 10.1101/2025.11.08.686969

**Authors:** Marcos E Jaso-Vera, Shohei Takaoka, Lanuza A.P. Faccioli, Zhiping Hu, Alejandro Soto-Gutierrez, Xiangbo Ruan

**Author notes:** **Corresponding**: Xiangbo Ruan, Ph.D. Assistant Professor of Medicine The Johns Hopkins University School of Medicine Division of Endocrinology, Diabetes and Metabolism Johns Hopkins All Children’s Hospital, Institute for Fundamental Biomedical Research 600 Fifth Street S. St. Petersburg, FL 33701.

## Abstract

Amino acid (AA) catabolism and ureagenesis in the liver are essential for maintaining systemic nitrogen homeostasis. Patients with metabolic-associated fatty liver disease (MAFLD) exhibit impaired hepatic AA catabolism and urea production, accompanied by repression of genes governing these pathways. The molecular basis of this repression remains unknown. Here, we identify the hepatocyte-specific long noncoding RNA hLMR1 as a key suppressor of hepatic AA catabolism. Using a humanized liver mouse model, we show that hLMR1 knockdown broadly upregulates genes involved in AA degradation and ureagenesis. Chromatin isolation by RNA purification (ChIRP) coupled with RNA-seq reveals extensive interactions between hLMR1 and the pre-mRNAs of AA catabolism genes, mediated by a conserved complementary motif within hLMR1 (nucleotides 587–598). In primary human hepatocytes, hLMR1 overexpression inhibits glucagon-induced activation of AA catabolic genes, whereas deletion of the 587–598 region abrogates this effect. Analysis of human liver RNA-seq datasets demonstrates a negative correlation between hLMR1 expression and AA catabolic gene programs in MAFLD. Together, these findings uncover hLMR1 as a previously unrecognized regulator of hepatic nitrogen metabolism, linking lncRNA dysregulation to metabolic dysfunction in MAFLD.

## Introduction

The liver plays a central role in maintaining whole-body nitrogen homeostasis. It is the primary site for catabolizing non-branched amino acids, including alanine, serine, glycine, glutamine, and histidine. Amino acid catabolism is tightly coupled to ureagenesis, a process that converts ammonia derived from amino acid deamination into urea for renal excretion^1^. In chronic metabolic diseases such as metabolic-associated fatty liver disease (MAFLD) and type 2 diabetes (T2D), hepatic capacity for amino acid catabolism and urea synthesis is markedly impaired^2–5^. Indeed, expression of key enzymes involved in nitrogen metabolism, including SDS, GLS2, CPS1, and ASS1, is significantly reduced in the livers of MAFLD patients^2, 6^. As a result, MAFLD patients often exhibit elevated circulating amino acid levels and hyperglucagonemia, likely driven by amino acid–stimulated glucagon release from pancreatic α-cells^7, 8^. This dysregulated liver–α-cell axis has emerged as a potential contributor to the pathogenesis of MAFLD and T2D^3, 4, 9^. However, the molecular mechanisms underlying the suppression of hepatic amino acid catabolism in humans remain poorly understood.

Long non-coding RNAs (lncRNAs) constitute a large class of RNA transcripts that exert regulatory control over diverse metabolic pathways^10–19^. Although lncRNA functions have been demonstrated in animal models, most human lncRNAs are poorly conserved, limiting mechanistic studies in mice^20^. To overcome this challenge, we employed a humanized liver mouse model^21^ in which human hepatocytes replace most mouse hepatocytes, allowing investigation of human-specific lncRNAs in vivo^14, 15^. Using this system, we previously identified a human-specific lncRNA, hLMR1 (human lncRNA metabolic regulator 1), which promotes hepatic cholesterol biosynthesis^15^. In the present study, RNA-seq analysis of humanized livers following hLMR1 knockdown revealed that hLMR1 represses a broad set of genes involved in amino acid catabolism and ureagenesis. Mechanistically, hLMR1 extensively interacts with the pre-mRNAs of amino acid catabolic genes, and this lncRNA–pre-mRNA interaction is essential for its repressive function. Consistent with this finding, analysis of human liver transcriptomes from MAFLD patients showed that hLMR1 expression negatively correlates with amino acid catabolism gene expression. Together, our results identify hLMR1 as a key suppressor of hepatic amino acid catabolism and suggest that its upregulation may contribute to nitrogen metabolic dysfunction in MAFLD.

## Results

### hLMR1 suppresses amino acid catabolism and ureagenesis in humanized liver mice

We previously identified hLMR1 as a non-conserved, liver-specific human lncRNA that is induced by feeding and promotes cholesterol synthesis^15^. Single-cell RNA-seq data from human liver tissues^22^ revealed that hLMR1, along with key amino acid (AA) catabolism genes, is highly enriched in hepatocytes (**Figure 1**). To systematically identify genes regulated by hLMR1, we performed high-throughput RNA sequencing on humanized mouse livers with or without hLMR1 knockdown. To specifically analyze the human transcriptome, RNA-seq reads were aligned to a combined human–mouse reference genome, and only human-specific reads were used for differential gene expression analysis (**Methods**).

**Figure 1.**
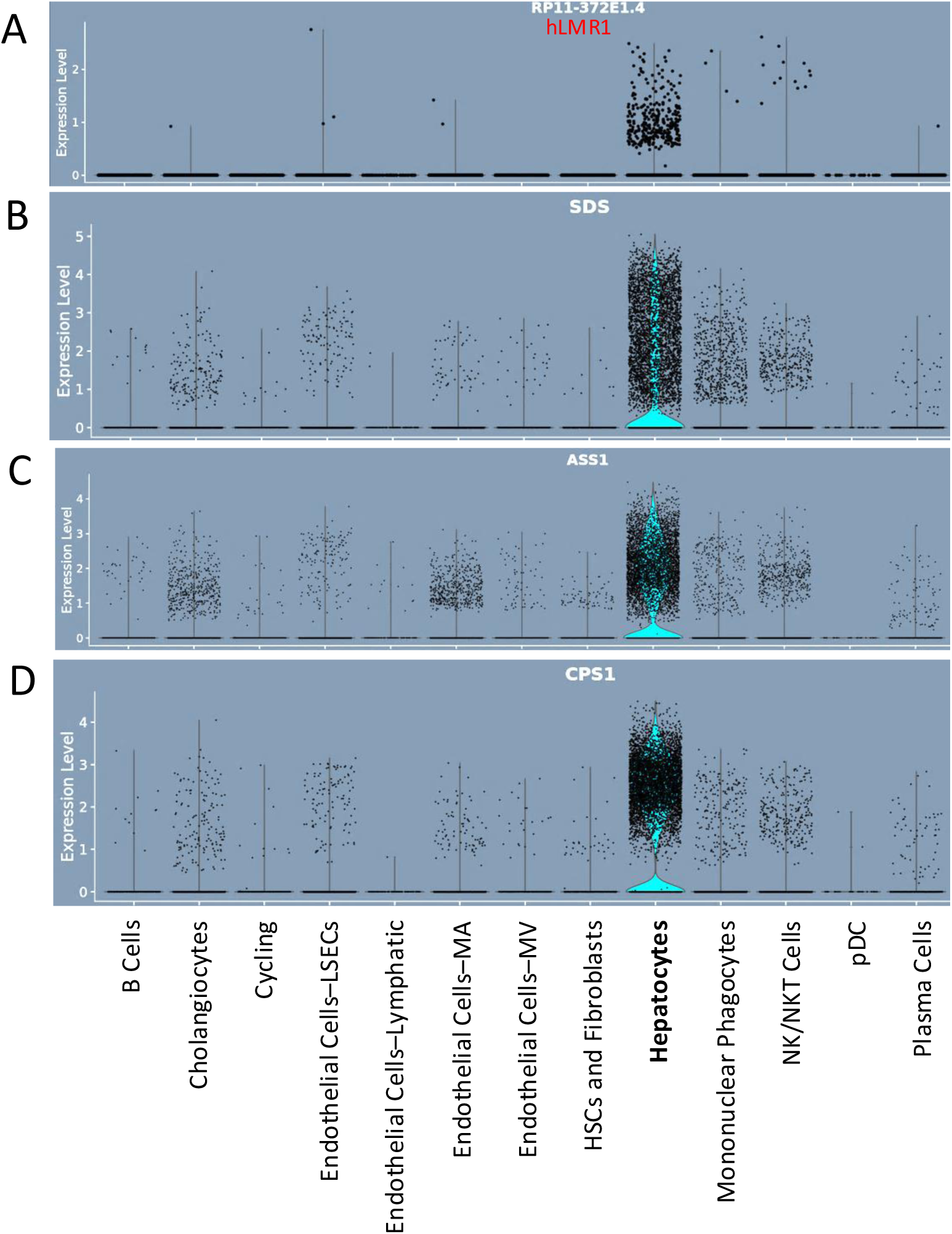
Expression of hLMR1 (**A**), SDS (**B**), ASS1 (**C**) and CPS1 (**D**) in major cell groups of human livers. Data were adapted from http://liveratlas-vilarinholab.med.yale.edu/.

As expected, hLMR1 knockdown reduced the expression of genes involved in cholesterol and fatty acid synthesis (**Supplementary Table 1**). Notably, hLMR1 depletion also led to robust upregulation of genes associated with amino acid metabolism and ureagenesis (**Table 1** and **Supplementary Table 1**). These included enzymes central to amino acid catabolism—such as serine dehydratase (SDS), glutaminase 2 (GLS2), and aminoadipate-semialdehyde synthase (AASS)—as well as urea cycle enzymes, including argininosuccinate synthase 1 (ASS1), carbamoyl-phosphate synthase 1 (CPS1), and arginase 1 (ARG1) (**Table 2**). Together, these data reveal that beyond promoting lipid synthesis, hLMR1 functions as a repressor of amino acid catabolism and ureagenesis in human hepatocytes.

**Table 1.** KEGG pathway analysis using the list of genes that were *upregulated* by knocking down of hLMR1 in humanized livers. Only Top 5 pathways were presented.

| KEGG_PATHWAY | P-Value | Benjamini | FDR |
| --- | --- | --- | --- |
| Biosynthesis of amino acids | 2.72E-05 | 3.56E-03 | 3.49E-03 |
| Arginine biosynthesis | 2.83E-05 | 3.56E-03 | 3.49E-03 |
| Complement and coagulation cascades | 8.69E-05 | 6.53E-03 | 6.40E-03 |
| Metabolic pathways | 1.04E-04 | 6.53E-03 | 6.40E-03 |
| Glycine, serine and threonine metabolism | 4.99E-04 | 2.50E-02 | 2.45E-02 |

**Table 2.**
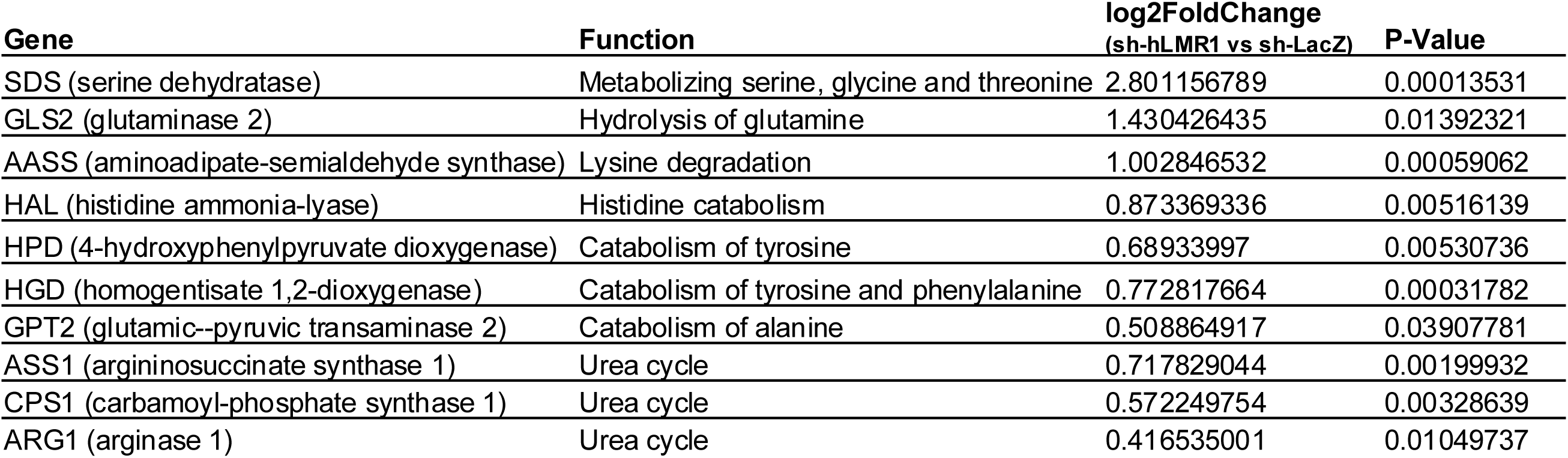
Representative genes in the AA catabolism and ureagenesis pathway that were *upregulated* by knocking down of hLMR1 in humanized livers.

| **Gene** | **Function** | **log2FoldChange**<br>**(sh-hLMR1 vs sh-LacZ)** | **P-Value** |
| --- | --- | --- | --- |
| SDS (serine dehydratase) | Metabolizing serine, glycine and threonine | 2.801156789 | 0.00013531 |
| GLS2 (glutaminase 2) | Hydrolysis of glutamine | 1.430426435 | 0.01392321 |
| AASS (aminoadipate-semialdehyde synthase) | Lysine degradation | 1.002846532 | 0.00059062 |
| HAL (histidine ammonia-lyase) | Histidine catabolism | 0.873369336 | 0.00516139 |
| HPD (4-hydroxyphenylpyruvate dioxygenase) | Catabolism of tyrosine | 0.68933997 | 0.00530736 |
| HGD (homogentisate 1,2-dioxygenase) | Catabolism of tyrosine and phenylalanine | 0.772817664 | 0.00031782 |
| GPT2 (glutamic--pyruvic transaminase 2) | Catabolism of alanine | 0.508864917 | 0.03907781 |
| ASS1 (argininosuccinate synthase 1) | Urea cycle | 0.717829044 | 0.00199932 |
| CPS1 (carbamoyl-phosphate synthase 1) | Urea cycle | 0.572249754 | 0.00328639 |
| ARG1 (arginase 1) | Urea cycle | 0.416535001 | 0.01049737 |

### hLMR1 interacts with pre-mRNAs of amino acid catabolism and ureagenesis genes

To explore the molecular basis of hLMR1-mediated repression, we performed Chromatin Isolation by RNA Purification (ChIRP)^23^ followed by RNA sequencing on human liver tissues. This approach enabled purification of endogenous hLMR1 and identification of RNA molecules interacting with it. To reduce interindividual variability, we pooled cryo-pulverized liver tissues from four male and four female donors. Two independent biotinylated oligo sets covering most of the hLMR1 transcript, and one control oligo set targeting the abundant lncRNA LINC01018^14^, were used. Real-time PCR confirmed specific enrichment of hLMR1 and LINC01018 by their corresponding probes (**Figure 2A**), validating the ChIRP assay. Only ChIRP signals overlapping between both hLMR1 oligo sets but absent in LINC01018 controls were defined as hLMR1-interacting RNA peaks (**Methods**).

**Figure 2.**
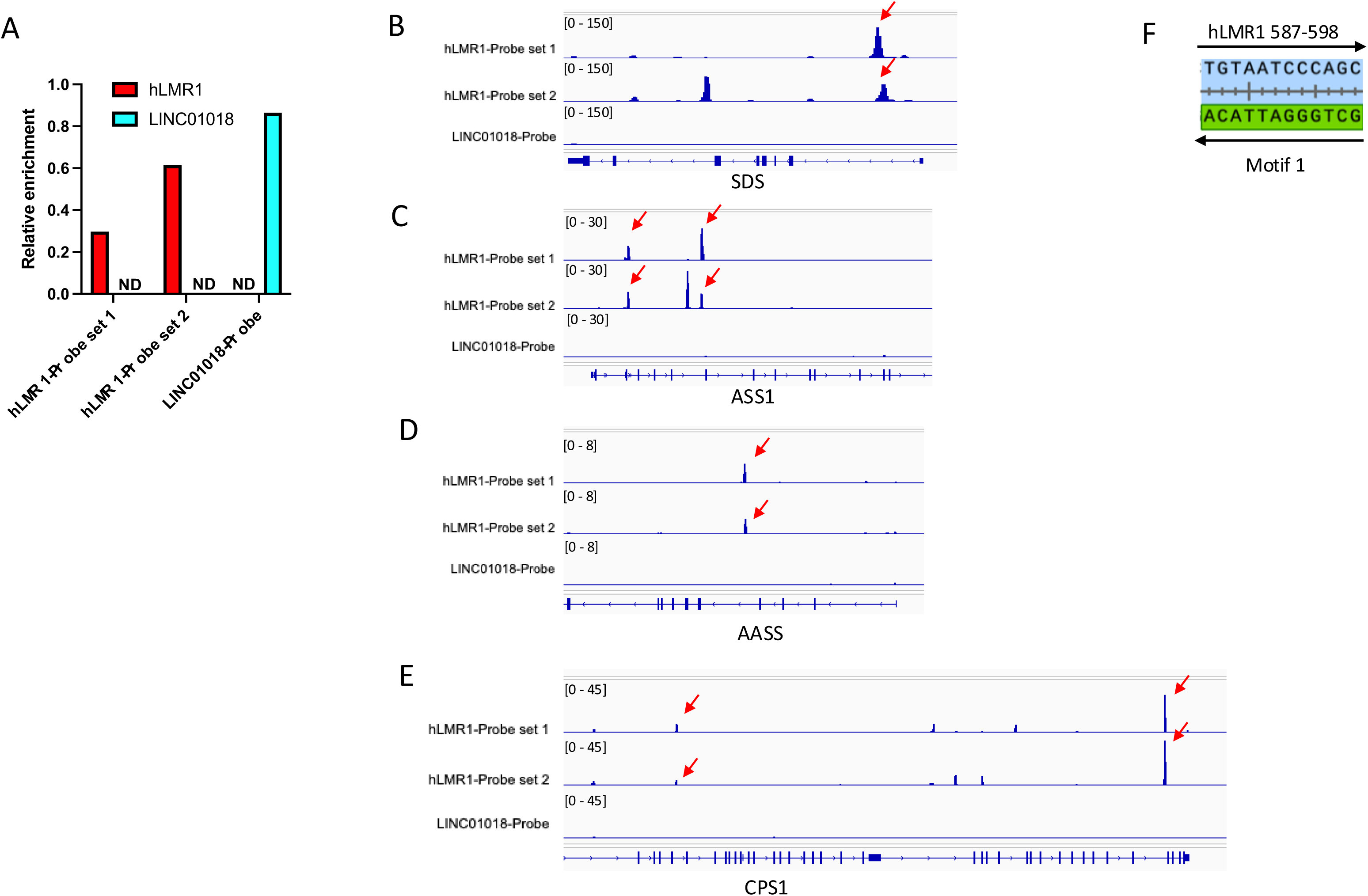
**A**. Relative enrichment of hLMR1 or LINC01018 in the ChIRP analysis. Relative enrichment is defined by the ratio between recovered portion and input. ND, not detected. **B-E**. Representative hLMR1 ChIRP RNA-seq peaks in SDS, ASS1, AASS, and CPS1. Peaks were visualized using CPM normalized bigwig files in IGV. Top overlapped peaks between hLMR1 probe sets were pointed by red arrows. ChIRP RNA-seq for another highly expressed lncRNA, LINC01018 was used as a negative control. All ChIRP experiments were performed using pooled human liver tissues. **F**. Illustration of hLMR1 sequence 587-598 matching with Motif 1.

Strong hLMR1 ChIRP signals were detected across the intronic regions of genes involved in amino acid catabolism and ureagenesis (**Figures 2B–2E**). KEGG pathway analysis of the top 1,000 hLMR1-interacting genes confirmed enrichment in amino acid metabolic pathways (**Table 3** and **Supplementary Table 2**). Motif analysis revealed several enriched RNA motifs (**Table 4**), among which the top motif (GCTGGGATTACA) was perfectly complementary to the hLMR1 sequence at nucleotides 587–598 (**Figure 2F**). These findings suggest that hLMR1 interacts directly with the pre-mRNAs of its target genes through a conserved complementary motif.

**Table 3.** KEGG pathway analysis using the list of top 1000 genes with peaks in hLMR1 ChIRP RNA-seq analysis.

| **KEGG_PATHWAY** | **P-Value** | **Benjamini** | **FDR** |
| --- | --- | --- | --- |
| Metabolic pathways | 1.91E-08 | 0.0000061 | 5.57E-06 |
| Complement and coagulation cascades | 7.54E-05 | 0.0121 | 0.011 |
| Cysteine and methionine metabolism | 0.000288 | 0.03 | 0.0274 |
| Fat digestion and absorption | 0.000375 | 0.03 | 0.0274 |
| Alanine, aspartate and glutamate metabolism | 0.000784 | 0.0377 | 0.0344 |

**Table 4.**
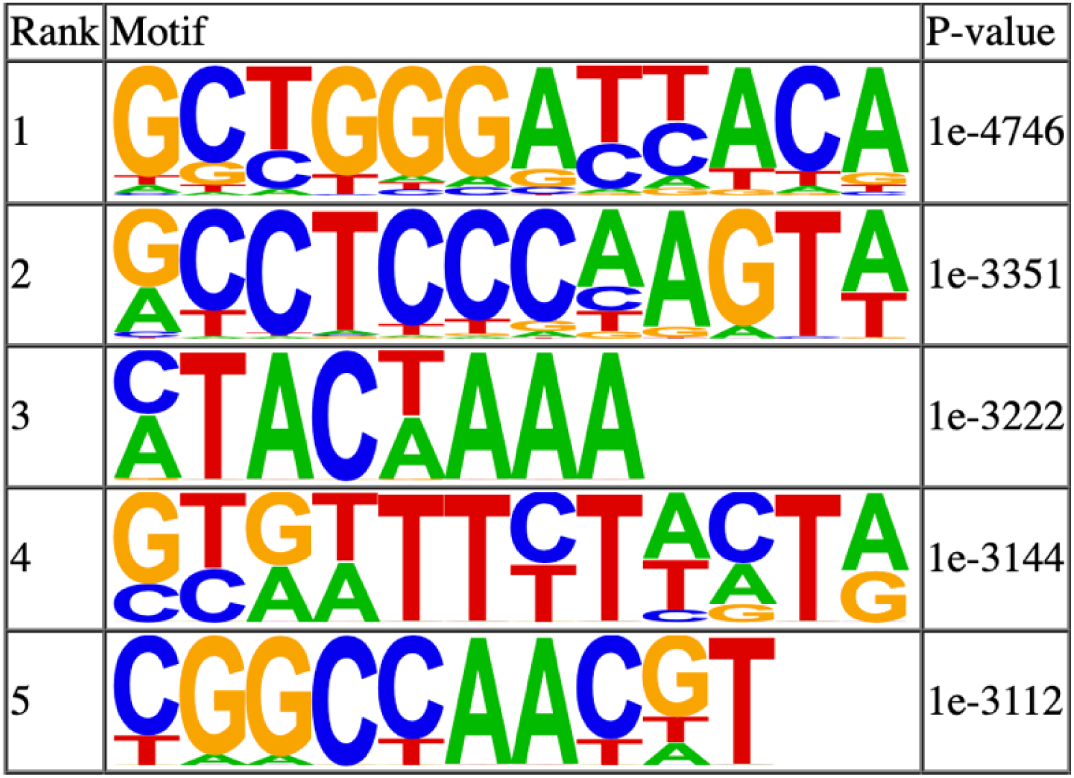
Top 5 motifs analyzed by HOMER (Hypergeometric Optimization of Motif EnRichment) using the sequences of hLMR1 ChIRP RNA-seq peaks.

### hLMR1 suppresses glucagon-induced amino acid catabolism in primary human hepatocytes

To test the functional significance of the hLMR1–pre-mRNA interaction, we generated adenoviruses expressing either wild-type hLMR1 or a mutant lacking the 587–598 motif (hLMR1Δ587–598). Primary human hepatocytes were infected with control, hLMR1, or hLMR1Δ587–598 adenoviruses and treated with glucagon for 10 hours. In control hepatocytes, glucagon robustly induced expression of SDS, CPS1, ASS1, HGD, and HAL, consistent with its known role in stimulating amino acid catabolism (**Figure 3**). Overexpression of hLMR1 completely blocked glucagon-induced SDS, CPS1, and HGD expression (**Figures 3C, 3E & 3G**), whereas induction of ASS1 and HAL was largely retained (**Figures 3B** & **3D**). In contrast, hLMR1Δ587–598 had no inhibitory effect, and glucagon induction of all five genes was comparable to control cells (**Figure 3**). Both constructs were expressed at similar levels (**Supplementary Figure 1**), excluding expression differences as a confounding factor.

**Figure 3.**
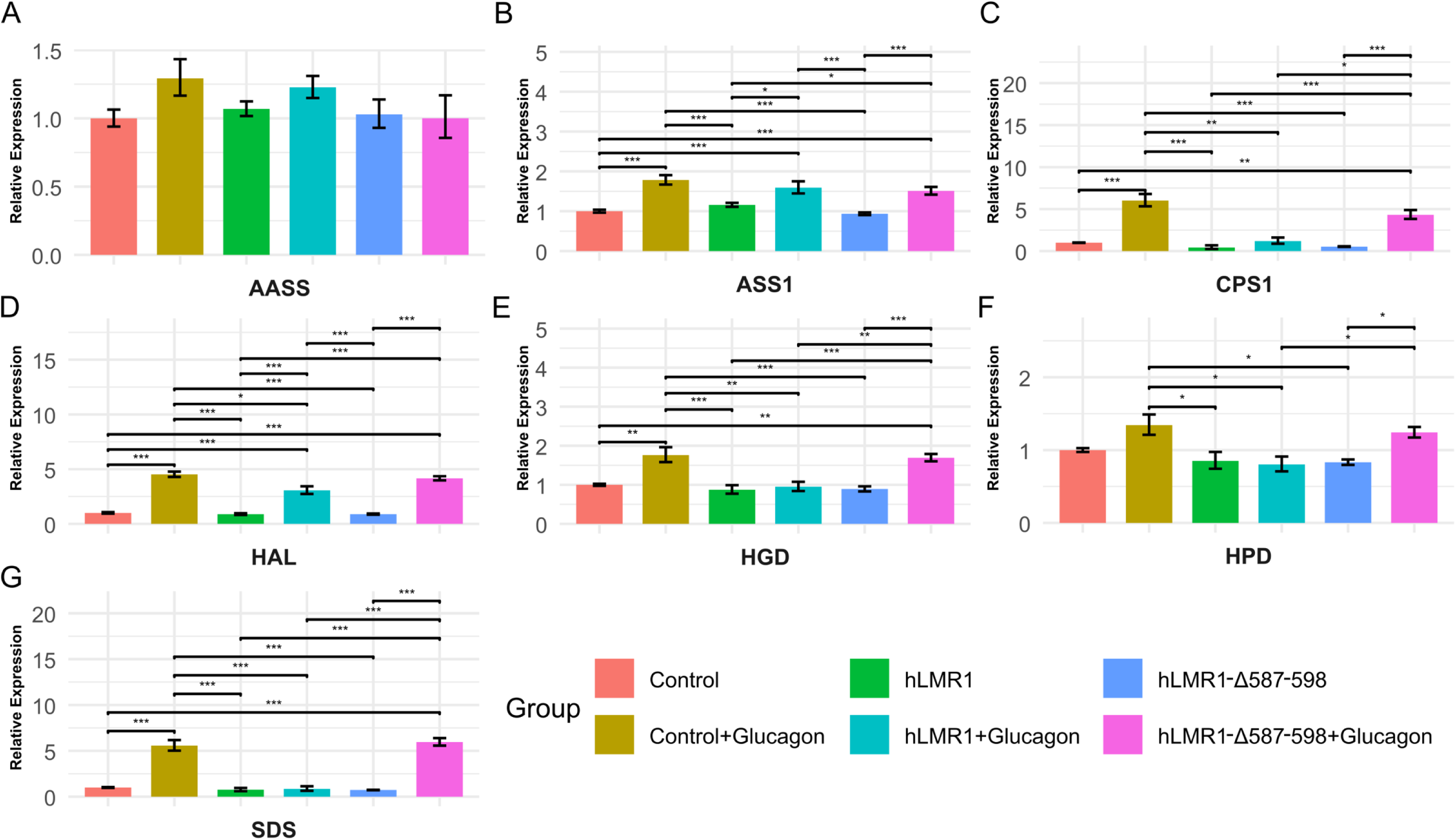
**A-G.** Relative mRNA expression of AASS, ASS1, CPS1, HAL, HGD, HPD and SDS in primary human hepatocytes (donor 1) infected with empty adenoviruses (Control) or adenoviruses expressing hLMR1, or hLMR1-1587-598. 36 hours after adenovirus infection, cells were treated with or without glucagon (50nM) for 10 hours and RNA were extracted for gene expression analysis. Data are shown as the geometric mean ± SEM. One-way ANOVA statistical analysis and Tukey HSD test were performed to calculate p-values. *, p < 0.05, **, p < 0.01, ***, p < 0.001.

Consistent with the transcriptional data, glucagon significantly increased urea production in control and hLMR1Δ587–598-expressing hepatocytes, but not in those overexpressing hLMR1 (**Figure 4**). These results indicate that hLMR1 suppresses glucagon-stimulated amino acid catabolism and ureagenesis likely through direct interaction with target pre-mRNAs.

**Figure 4.**
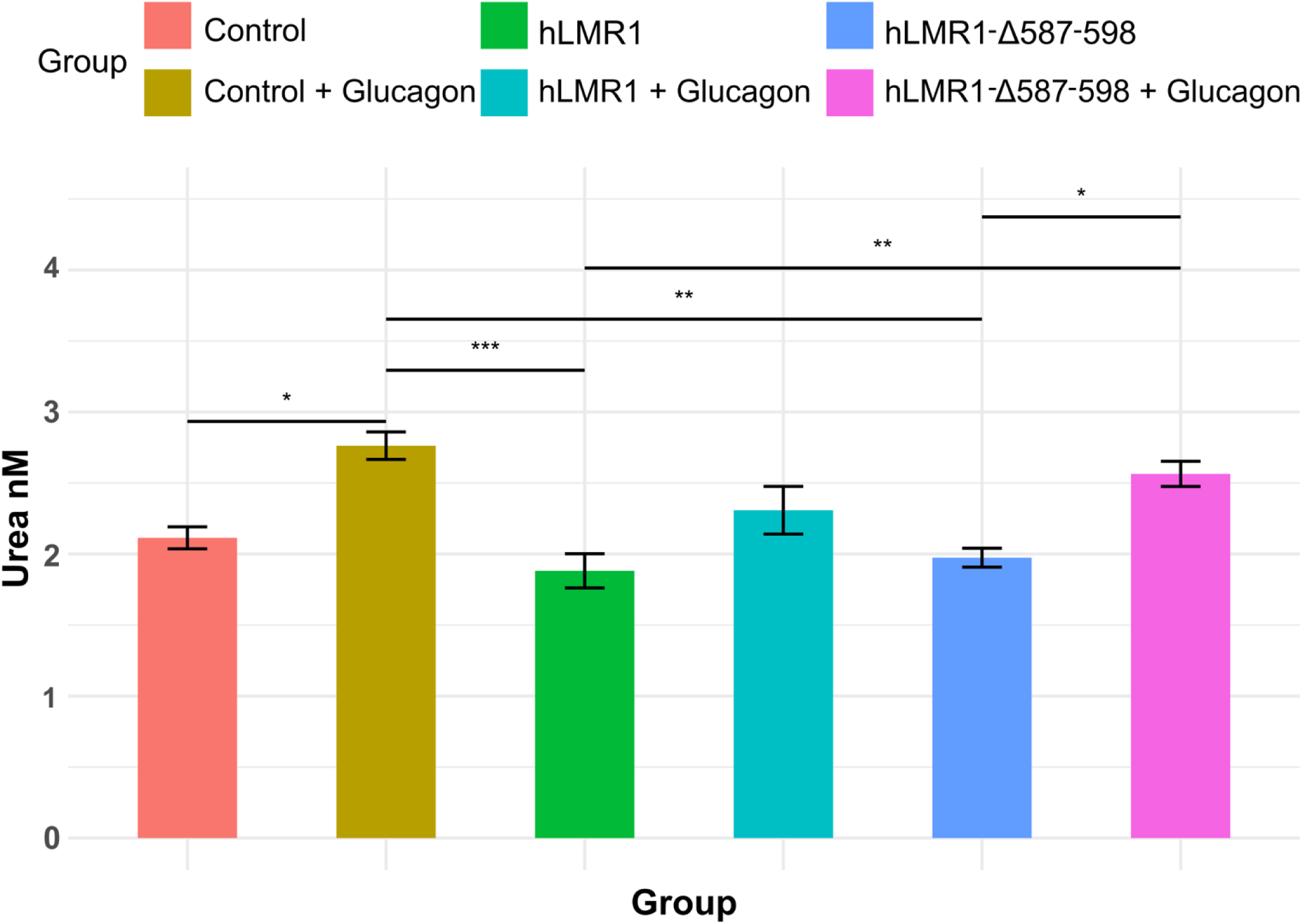
Urea levels in the cell culture medium of cultured primary human hepatocytes as in Figure 3. Data are shown as the geometric mean ± SEM. One-way ANOVA statistical analysis and Tukey HSD test were performed to calculate p-values. *, p < 0.05.

To exclude the possibility that our observation is limited to a specific donor, we performed the experiment in human primary hepatocytes from a second donor. As shown in **Figure 5**, we observed a similar result showing glucagon robustly induced AA catabolism genes in control hepatocytes and hepatocytes expressing hLMR1-Δ587-598. In contrast, glucagon-induced expression of SDS and HGD was significantly supressed (**Figures 5E & 5G**), and a trend of suppression for ASS1, CPS1, and HAL (**Figures 5B, 5C & 5D**) was observed in hepatocytes overexpressing hLMR1. We also found the overexpression of hLMR1 and hLMR1-Δ587-598 is comparable in this setting (**Supplementary Figure 2**). These results support that the role of hLMR1 in suppressing AA catabolism is not limited to a specific donor or genetic background.

**Figure 5.**
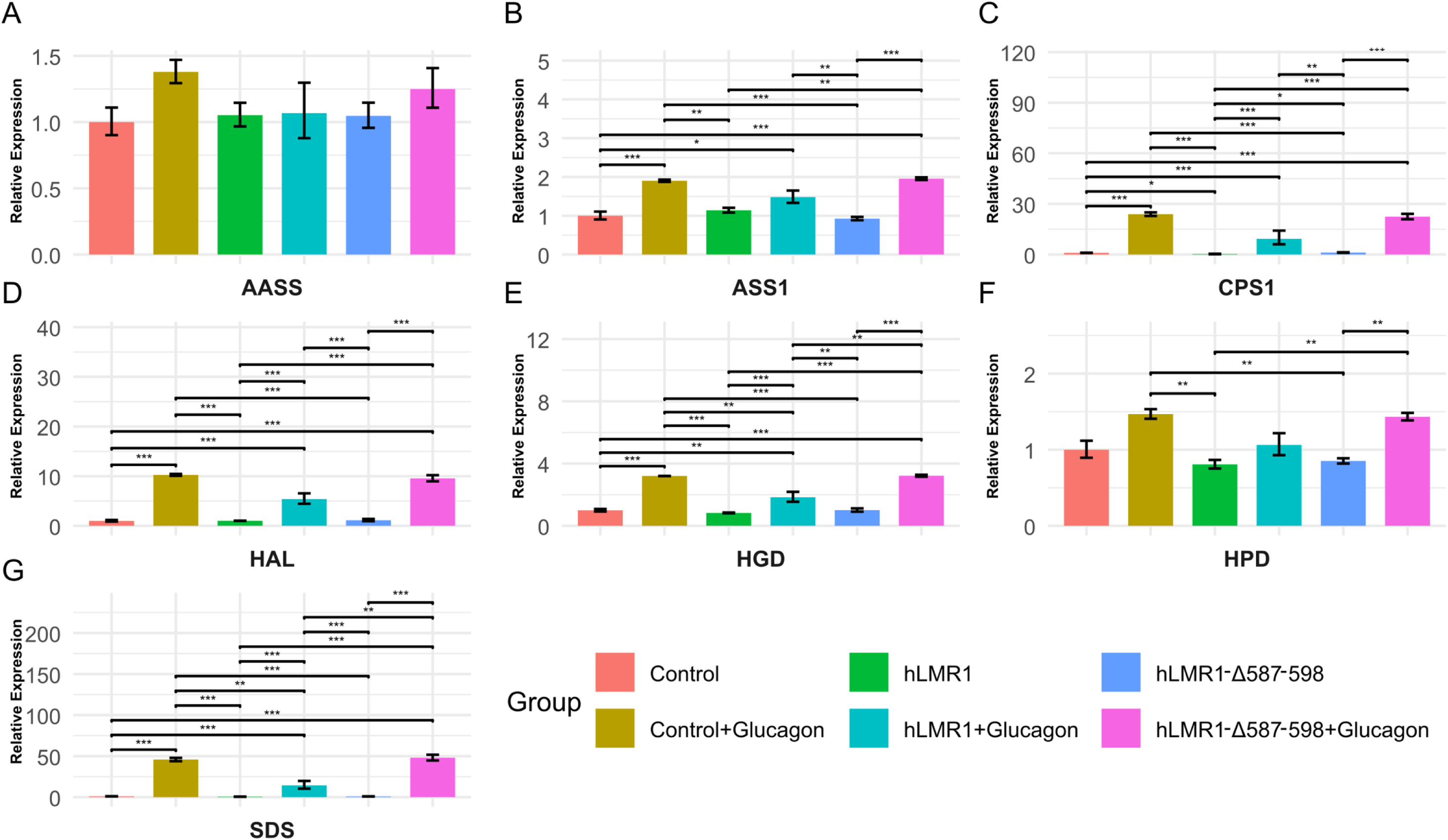
**A-G.** Relative mRNA expression of AASS, ASS1, CPS1, HAL, HGD, HPD and SDS in primary human hepatocytes (donor 2) infected with empty adenoviruses (Control) or adenoviruses expressing hLMR1, or hLMR1-1587-598. 36 hours after adenovirus infection, cells were treated with or without glucagon 50nM for 10 hours and RNA were extracted for gene expression analysis. Data are shown as the geometric mean ± SEM. One-way ANOVA statistical analysis and Tukey HSD test were performed to calculate p-values. *, p < 0.05, **, p < 0.01, ***, p < 0.001.

### hLMR1 does not repress amino acid catabolism in mouse hepatocytes

Given that hLMR1 is a human-specific lncRNA, we next examined whether its overexpression could similarly suppress amino acid catabolism in mouse hepatocytes. Primary mouse hepatocytes were infected with adenoviruses expressing hLMR1 or hLMR1Δ587–598 and treated with glucagon as above. Despite efficient transgene expression (**Supplementary Figure 3**), neither construct altered glucagon-induced activation of amino acid catabolism or ureagenesis genes (**Figure 6**). These findings confirm that the repressive function of hLMR1 is species-specific.

**Figure 6.**
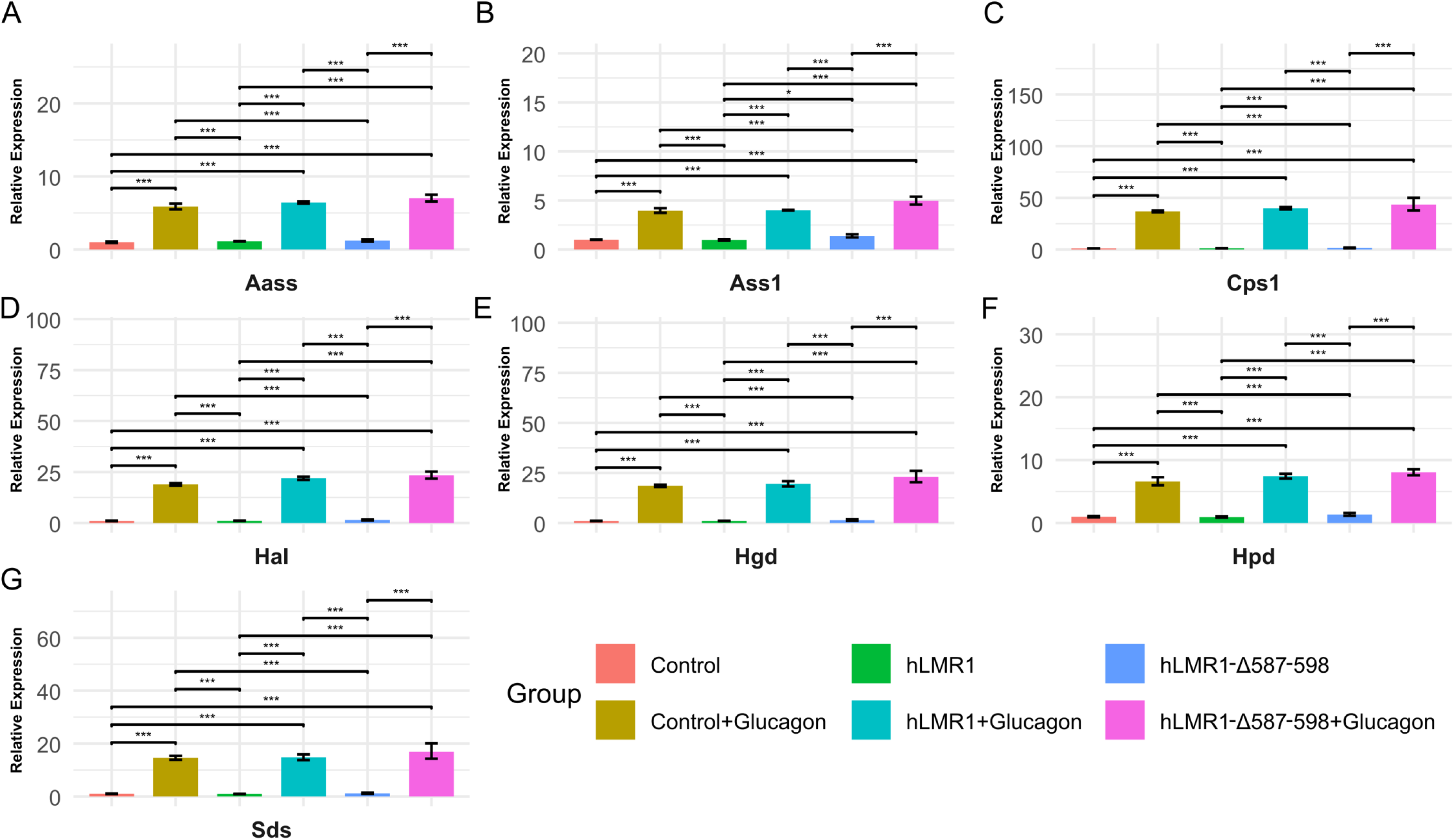
**A-G.** Relative mRNA expression of Aass, Ass1, Cps1, Hal, Hgd, Hpd and Sds in primary mouse hepatocytes infected with empty adenoviruses (Control) or adenoviruses expressing hLMR1, or hLMR1-1587-598. 36 hours after adenovirus infection, cells were treated with or without glucagon 10nM for 10 hours and RNA were extracted for gene expression analysis. Data are shown as the geometric mean ± SEM. One-way ANOVA statistical analysis and Tukey HSD test were performed to calculate p-values. *, p < 0.05, **, p < 0.01, ***, p < 0.001.

### Hepatic hLMR1 expression inversely correlates with amino acid catabolism genes in MAFLD

To evaluate the physiological relevance of hLMR1, we analyzed a large RNA-seq dataset from liver biopsies across the MAFLD spectrum^24^. hLMR1 expression increased from healthy controls to MASH (**Figure 7**). In contrast, representative amino acid catabolism and ureagenesis genes, including SDS and ASS1, were reciprocally downregulated. In an independent interventional study^25^, one week of low-carbohydrate dietary treatment reduced hepatic hLMR1 expression while increasing SDS, GLS2, AASS, ASS1, and CPS1 expression in obese MAFLD patients (**Table 5**). These clinical observations support that elevated hLMR1 may contribute to impaired amino acid catabolism and ureagenesis in MAFLD.

**Figure 7.**
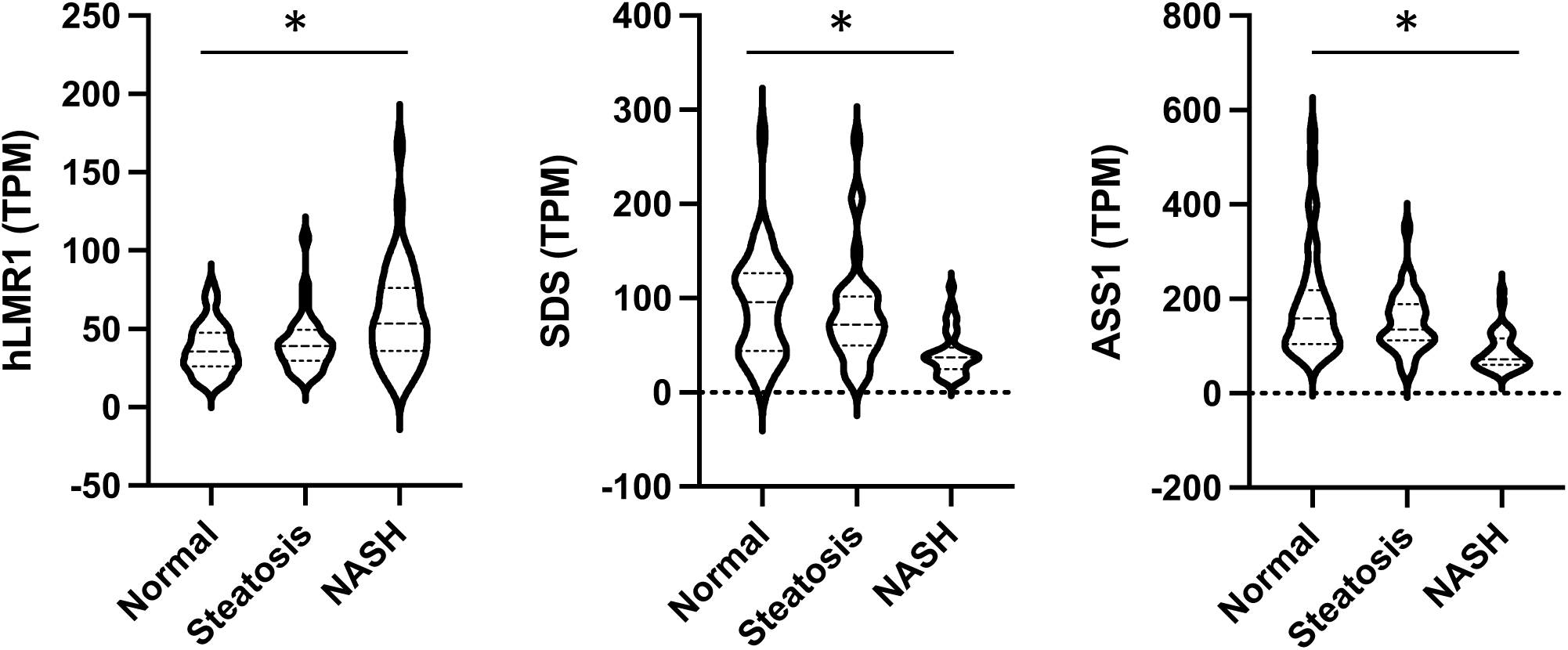
Relative RNA expression of hLMR1, SDS and ASS1 in human liver tissue samples from health individuals (Normal), individuals with hepatic steatosis, and MASH. TPM, transcript per million. * p<0.05, by one-way ANOVA analysis. Data were analyzed from raw data of NCBI BioProject: PRJNA512027.

**Table 5.** Hepatic expression of hLMR1 and representative amino acid catabolism genes in obese individuals with NAFLD before and after a low carb diet intervention. Data were reanalyzed from NCBI dataset GSE107650 and a paired t test was performed to calculate the p value.

| Gene | Log2(fold_change)<br>After vs Before | P-Value |
| --- | --- | --- |
| hLMR1 | -0.733363659 | 7.9371E-05 |
| SDS | 0.692498252 | 0.01168179 |
| GLS2 | 1.127643149 | 3.1486E-16 |
| AASS | 0.730033593 | 0.00243507 |
| ASS1 | 1.045257926 | 3.3781E-12 |
| CPS1 | 1.421943193 | 2.2768E-06 |

## Discussion

Maintaining systemic nitrogen homeostasis is a fundamental function of the liver. Hepatocytes catabolize major non-branched amino acids, including alanine, glutamine, and serine, and couple this process with ureagenesis to safely eliminate ammonia from circulation. However, hepatic capacity for amino acid catabolism and urea synthesis is markedly reduced in metabolic diseases such as metabolic-associated fatty liver disease (MAFLD) and type 2 diabetes (T2D)^2–7, 9, 26^. Recent studies have highlighted disruption of the liver–α-cell axis as a shared feature of both disorders, yet the molecular mechanisms underlying the loss of glucagon-driven amino acid catabolism remain unclear. Here, we identify the human hepatocyte-specific long noncoding RNA (lncRNA), hLMR1, as a key regulator of this pathway. Our findings, together with clinical data, suggest that upregulation of hLMR1 contributes to impaired amino acid catabolism and ureagenesis in MAFLD.

Multiple lines of evidence support this conclusion. First, hLMR1 is highly and specifically expressed in hepatocytes (**Figure 1**), consistent with the hepatocyte-specific nature of amino acid catabolism and ureagenesis. Second, under physiological conditions, hLMR1 expression is suppressed by fasting, whereas amino acid catabolism genes are strongly induced, suggesting reciprocal regulation^14, 15^. Third, hLMR1 is predominantly localized in the nucleus of human hepatocytes^15^, consistent with its proposed mechanism of interacting with the pre-mRNAs of amino acid catabolism genes to repress their transcription. Fourth, in human NAFLD patients subjected to a low-carbohydrate dietary intervention^25^, hepatic hLMR1 expression was reduced, while amino acid catabolism genes were upregulated (**Table 5**)—aligning with our prior observation that glucose availability promotes hLMR1 expression^27^.

Our mechanistic data support a model in which hLMR1 directly interacts with the pre-mRNAs of amino acid catabolism genes to suppress their expression. Among the most strongly regulated targets is SDS, whose first intron contains several hLMR1-interacting motifs. Similar lncRNA–pre-mRNA regulatory mechanisms have been described previously, where lncRNAs act as organizing hubs to coordinate the expression of functionally related genes^28^. Our results extend this concept to hepatic nitrogen metabolism: amino acid catabolism and ureagenesis genes are highly expressed during fasting, when glucagon-driven transcription predominates, and must be efficiently silenced upon feeding. We propose that feeding-induced hLMR1 acts as a nuclear scaffold that binds pre-mRNAs of these genes to repress their expression, thereby enabling rapid metabolic switching between fasting and fed states.

Consistent with its species specificity, hLMR1 failed to suppress glucagon-induced amino acid catabolism in primary mouse hepatocytes, indicating that its regulatory function is unique to humans (**Figure 6**). Whether a murine homolog exists or alternative mechanisms fulfill an equivalent role in mice remains unknown, suggesting an additional regulatory layer for nitrogen metabolism that evolved specifically in humans.

Several important questions remain. As hLMR1 regulates both lipid and amino acid metabolic pathways, future studies should define the precise RNA–RNA and RNA– protein interactions underlying its function. Mapping hLMR1’s structural domains and testing how specific mutations affect target recognition will be essential to determine whether secondary RNA folding contributes to its regulatory activity. Another key direction is to assess whether inhibition of hLMR1 can restore the impaired liver–α-cell axis observed in MAFLD and T2D^4, 9, 26^. Given that hLMR1 promotes hepatic cholesterol synthesis and is expressed exclusively in hepatocytes, pharmacological suppression of hLMR1—using antisense oligonucleotides or GalNAc-conjugated siRNAs—could simultaneously ameliorate hyperlipidemia and hyperglycemia. Because hLMR1 is human-specific, such therapeutic strategies can only be evaluated in diet-induced MAFLD/T2D models in humanized liver mice^29–31^ before clinical studies.

## Methods

### Bulk RNA-seq Analysis of Humanized Liver Mouse Data

Bulk RNA-seq samples were processed by the Illumina stranded total RNA ligation prep kit and sequenced on the Illumina NovaSeq S1 200 platform at the The Single Cell & Transcriptomics Core of Johns Hopkins School of Medicine. The quality of raw sequencing reads (FASTQ files) was assessed using FastQC (version 0.11.8). Adapter sequences and low-quality reads were trimmed or removed using Trimmomatic (version 0.39)^32^. Prior to alignment, the GRCh38 Ensembl human genome and the GRCm39 Ensembl mouse genome were merged to enable host–graft read discrimination. Filtered reads were aligned to the merged genome using STAR (version 2.7.8a)^33^. Uniquely mapped human reads were extracted from the resulting BAM files using SAMtools (version 1.13)^34^. Gene-level read counts for mature mRNA were generated with featureCounts from the Subread package (version 2.0.0)^35^. Genes expressed at raw counts greater than one in two or fewer samples were excluded prior to normalization.

The count matrix was normalized and analyzed for differential expression using DESeq2 (version 1.36.0) [PMID: 25516281] in R (version 4.1.0). Due to high biological variability among humanized liver samples, differentially expressed genes (DEGs) were defined as those with P ≤ 0.05 and an absolute log2 fold change (|Log2FC|) ≥ 0.38. DEGs with a baseMean among the top 10,000 were used for KEGG pathway enrichment analysis using the DAVID functional annotation tool^36^.

### Chromatin Isolation by RNA Purification (ChIRP)

The ChIRP protocol in this study is based on the previous publication^37^ with slight modifications. Pooled human liver tissues from eight donors (4 male and 4 female) were minced into thin slices and crosslinked in 3% formaldehyde (1 mL per 50 mg tissue) (Thermo Cat#28906) for 30 minutes at room temperature. Crosslinking was quenched by adding 0.125 mM glycine (CST Cat#7005S) and incubating for 5 minutes. The tissue was pelleted at 2,000 × g, washed once with PBS, pelleted again, and the supernatant carefully aspirated. The pellet was resuspended in cell lysis buffer (100 mg tissue per 1 mL buffer, 50 mM Tris-HCl pH 7.0, 10 mM EDTA, and 1% SDS). Cells were disrupted using a tissue grinder to release crosslinked RNA, DNA, and protein complexes.

Chromatin was sheared by sonication using a Bioruptor (Diagenode Cat#B01020001) for 1 hour. To confirm appropriate shearing, a 5 µL aliquot was purified using a PCR purification kit and analyzed by agarose gel electrophoresis (Qiagen Cat#28106), ensuring an average fragment size below 500 bp. The lysate was clarified by centrifugation at 16,000 × g for 10 minutes at 4 °C, and the supernatant was transferred to fresh tubes.

For hybridization, samples were diluted threefold in hybridization buffer (750 mM NaCl, 1% SDS, 50 mM Tris-HCl pH 7.0, 1 mM EDTA, and 15% formamide) and incubated with biotinylated RNA-specific ChIRP probes (1 µL of 100 µM probe mix per 1 mL lysate). Two independent probe libraries targeting unique regions of the hLMR1 transcript were used (**Supplementary Table 3**). A probe library targeting LINC01018 served as a negative control. Hybridization was carried out at 37 °C for 16 hours with end-to-end rotation.

After hybridization, 100 µL of C-1 Dynabeads (Invitrogen Cat#65001) were added per 1 µL of 100 µM probes and incubated at 37 °C for 30 minutes. Beads were then washed five times with wash buffer and transferred to a new tube for RNA extraction. RNase A-treated samples were included as negative controls, and input lysates served as positive controls.

To recover RNA, bead-bound and input samples were treated with proteinase K (5 µL in 95 µL RNA PK buffer, 100 mM NaCl, 10 mM Tris-HCl pH 8.0, 1 mM EDTA, and 0.5% SDS) for 45 minutes at 50 °C with end-to-end shaking, followed by heat inactivation at 95 °C for 10 minutes. RNA was then extracted using Trizol–chloroform, and the aqueous phase was purified using RNeasy Mini columns following the manufacturer’s clean-up and on-column DNase digestion protocols (Qiagen Cat#74106). Purified RNA was eluted in nuclease-free water and used for cDNA synthesis. ChIRP efficiency was evaluated by qPCR using oligonucleotides specific to hLMR1 and LINC01018 (negative control).

### Chromatin Isolation by RNA Purification (ChIRP) Followed by RNA-seq Analysis

ChIRP-RNA samples were processed by the Takara SMARTer stranded v3 total RNA library prep kit and sequenced on the Illumina NovaSeq S1 200 platform at the The Single Cell & Transcriptomics Core of Johns Hopkins School of Medicine. Sequencing read quality was assessed using FastQC (version 0.11.8), and adapter sequences and low-quality reads were removed with fastp (version 0.23.2)^38^. Filtered reads were aligned to the GRCh38 Ensembl human genome using STAR (version 2.7.8a). Reads with mapping quality scores below 30 were excluded using SAMtools (version 1.13). PCR duplicates were removed with Picard (version 2.27.4) [RRID: SCR_006525].

ChIRP-enriched RNA peaks were identified using MACS2 (version 2.2.7.1)^39^ in single-end mode with matched input controls. To optimize peak detection, the mfold lower and upper parameters were adjusted to 2 and 100, respectively (default values: 5 and 50). Peaks overlapping ENCODE Blacklist regions for hg38 were excluded using bedtools (version 2.24.0)^40^. Remaining peaks were annotated using the annotatePeaks.pl command from HOMER (version 4.11)^41^.

Only peaks overlapping between two independent probe sets targeting hLMR1, while not overlapping with peaks derived from LINC01018-targeting oligos, were retained using bedtools filtering. The resulting high-confidence hLMR1-interacting RNA peaks were analyzed for regulatory motif enrichment with the findMotifsGenome.pl command from HOMER. The top 1000 filtered peaks, ranked by MACS2 peak score, were further used for KEGG pathway enrichment analysis using DAVID^36^.

### Adenovirus Construction, Production, and Titration

Full-length hLMR1 and deletion mutant hLMR1 Δ587–598 sequences were cloned into the pAd/CMV/V5-DEST vector using the Gateway cloning system (Invitrogen). PCR amplification was used to introduce attB recombination sites into the hLMR1 sequence derived from pCDNA3.1 plasmids (**Supplementary Table 3**). The amplified DNA fragments were purified from a 1% agarose gel and cloned into the pDONR221 entry vector using the BP recombination reaction according to the manufacturer’s instructions (Invitrogen Cat#12536017 and Cat#11789021). LR recombination was then performed to transfer the inserts into the pAd/CMV/V5-DEST destination vector (Invitrogen Cat#V49320). The integrity of all plasmids was verified by whole-plasmid sequencing.

For adenovirus production, six 15-cm dishes of 293A cells were transfected with 2 µg of PacI-linearized pAd/CMV/V5-DEST–hLMR1 or Δ587–598 plasmids using Lipofectamine 2000 (Thermo Fisher Scientific, Cat#11668019), following the manufacturer’s protocol. Two days post-transfection, cells were transferred to 10-cm dishes, and the medium was replaced every two days until a clear cytopathic effect was observed under the microscope. When more than 80% of the cells had detached, both cells and medium were collected, and three freeze–thaw cycles were performed to release the crude virus.

For virus amplification and purification, 293A cells were infected with the crude viral lysate and incubated for 48 hours. Cells were harvested, resuspended in PBS, and subjected to three additional freeze–thaw cycles to release viral particles. The lysate was centrifuged to remove cell debris, and the supernatant was purified using a 50% cesium chloride (CsCl) gradient (32,000 × g, 4 °C, 18 h). The viral band was collected from the top of the gradient using an 18-gauge syringe, desalted by gravity flow through a PD-10 desalting column (GE Healthcare), and quantified by measuring OD260/280 with a Cytation spectrophotometer. The virus was diluted to a final concentration of 1 × 10¹² particles/mL and stored at −80 °C until use.

Virus titration was performed using the Adeno-X Rapid Titer Kit (Takara Bio, Cat#632250). Briefly, 293A cells were seeded in 24-well plates and infected with serial dilutions (10⁻², 10⁻⁴, 10⁻⁵, 10⁻⁶, and 10⁻⁷) of the purified virus. After 48 hours of incubation, cells were fixed with ice-cold methanol and washed three times with PBS containing 1% BSA. An anti-hexon antibody was applied and incubated for 1 hour at 37 °C, followed by three PBS washes. A horseradish peroxidase (HRP)-conjugated rat anti-mouse secondary antibody was then applied for 1 hour at 37 °C. After washing three times with PBS containing 1% BSA, a DAB working solution was added to visualize infection foci. Black-stained foci were counted at 20× magnification, and infectious units (IFU/mL) were calculated for each dilution. The average IFU/mL across dilutions was used as the final virus titer, which was applied in subsequent experiments.

### Primary Human Hepatocyte (PHH) Culture

Diseased adult human liver cells were obtained from the Human Organ processing hub and from the Center for Transcriptional Medicine at the University of Pittsburgh. The Institutional Review Board at the University of Pittsburgh has approved our protocol and given the Not Human Research Determination (IRB# STUDY24020093). Hepatocytes were isolated using a three-step collagenase digestion technique as previously described^42^. Cells were kept on ice throughout handling until plating. Cell suspensions were gently resuspended and diluted fourfold with ice-cold Williams’ E Medium (WEM) (Thermo Fisher Scientific, Cat#A1217601). The diluted cells were centrifuged at 600 rpm for 7 minutes at 4 °C, and the pellet was resuspended in ice-cold WEM followed by a second centrifugation at 500 rpm for 5 minutes at 4 °C. After washing, the cell pellet was resuspended in 10 mL of ice-cold WEM supplemented with 5% FBS. Viable cells were counted using a Neubauer hemocytometer with trypan blue exclusion. Cells were then diluted in WEM containing 10% FBS at 37 °C to a final concentration of 1 × 10⁶ cells/mL. A total of 0.5 mL of the suspension was plated per well in a 24-well plate pre-coated with collagen. After 3 hours of attachment, cells were washed twice with PBS, and pre-warmed WEM containing the designated treatments was added. Cultures were maintained at 37 °C in 5% CO_₂_.

### Primary Mouse Hepatocyte (PMH) Culture

Primary mouse hepatocytes were isolated from C57BL/6J mice by in situ perfusion. The liver was perfused via the vena cava with 30 mL of pre-warmed perfusion buffer (HBSS containing 0.5 mM EDTA, pH 8.0, and 25 mM HEPES, pH 7.5) at a flow rate of 1 mL/min. Immediately afterward, 30 mL of pre-warmed digestion buffer (HBSS containing 25 mM HEPES, pH 7.5, and 1 mg/mL Collagenase D) was perfused at the same rate. The digested liver was excised and placed in ice-cold DMEM supplemented with 10% FBS. Hepatocytes were gently released by teasing the liver capsule with forceps, and the resulting cell suspension was filtered through a 100 µm cell strainer. Cells were pelleted by centrifugation at 50 × g for 5 minutes at 4 °C, resuspended in fresh DMEM + 10% FBS, and washed twice by repeating the centrifugation step. The final pellet was resuspended in 5 mL of DMEM + 10% FBS and kept on ice. Cell number and viability were determined using trypan blue exclusion. Cells were diluted in WEM + 10% FBS at 37 °C to a final concentration of 1 × 10^⁶^ cells/mL, and 0.5 mL of the suspension was added to each well of a collagen-coated 24-well plate. After 3 hours of attachment, cells were washed twice with PBS and incubated in pre-warmed WEM containing the required treatments. Cultures were maintained at 37 °C in 5% CO_₂_.

### Primary Hepatocyte Transduction and Treatment

For viral transduction, adenoviral vectors were added to WEM containing 10% FBS at a multiplicity of infection (MOI) of 20. The virus-containing medium was added to the hepatocyte cultures and incubated for 24 hours. Cells were then washed twice with PBS and replenished with fresh sterile WEM + 10% FBS, followed by overnight incubation under the same conditions. The next morning, PHH were washed twice with PBS and incubated with WEM containing 2.5 mM glucose for 3 hours. Cells were then washed twice with PBS and treated with either WEM containing 2.5 mM glucose or WEM supplemented with 50 nM glucagon and 2 mM pyruvate, as described in the text. PHH were incubated for 11 hours, after which the culture medium was collected and stored at −80 °C. Cells were washed once with ice-cold PBS and lysed in RLT buffer for RNA extraction using the RNeasy Mini Kit.

### Complementary DNA Synthesis and Quantitative PCR (RT-qPCR)

Total RNA was extracted using the RNeasy Mini Kit with on-column DNase digestion. RNA was eluted in nuclease-free water, and 500 ng of RNA per sample was used for cDNA synthesis with SuperScript III Reverse Transcriptase (Thermo Fisher, Cat#11752050). The resulting cDNA was diluted 1:10 in nuclease-free water. Quantitative PCR was performed using SYBR Green reagents on a QuantStudio 7 instrument. Primer sequences used for qPCR are listed in (**Supplementary Table 3**).

### Urea Quantitation in Cell Culture Media

Culture media from PHH experiments were thawed on ice and directly analyzed for urea content using the Urea Assay Kit (Abcam, Cat# ab83362), following the manufacturer’s instructions. Absorbance was measured using a Cytation microplate reader, and urea concentrations were calculated according to the standard curve.

### Statistics

Statistical analysis was performed using R version 4.5.0. Student’s t-test was used for comparisons between two groups; One-way ANOVA was used for comparisons between three or more groups, followed by post hoc analysis with Tukey-HSD. ANOVA p-values reported in each bar chart represent the overall significance of the differences among the groups being compared. Significant differences among each individual group were annotated by line illustrations and stars, which were automatically generated in R.

## Figure Legend

**Supplementary Figure 1.**
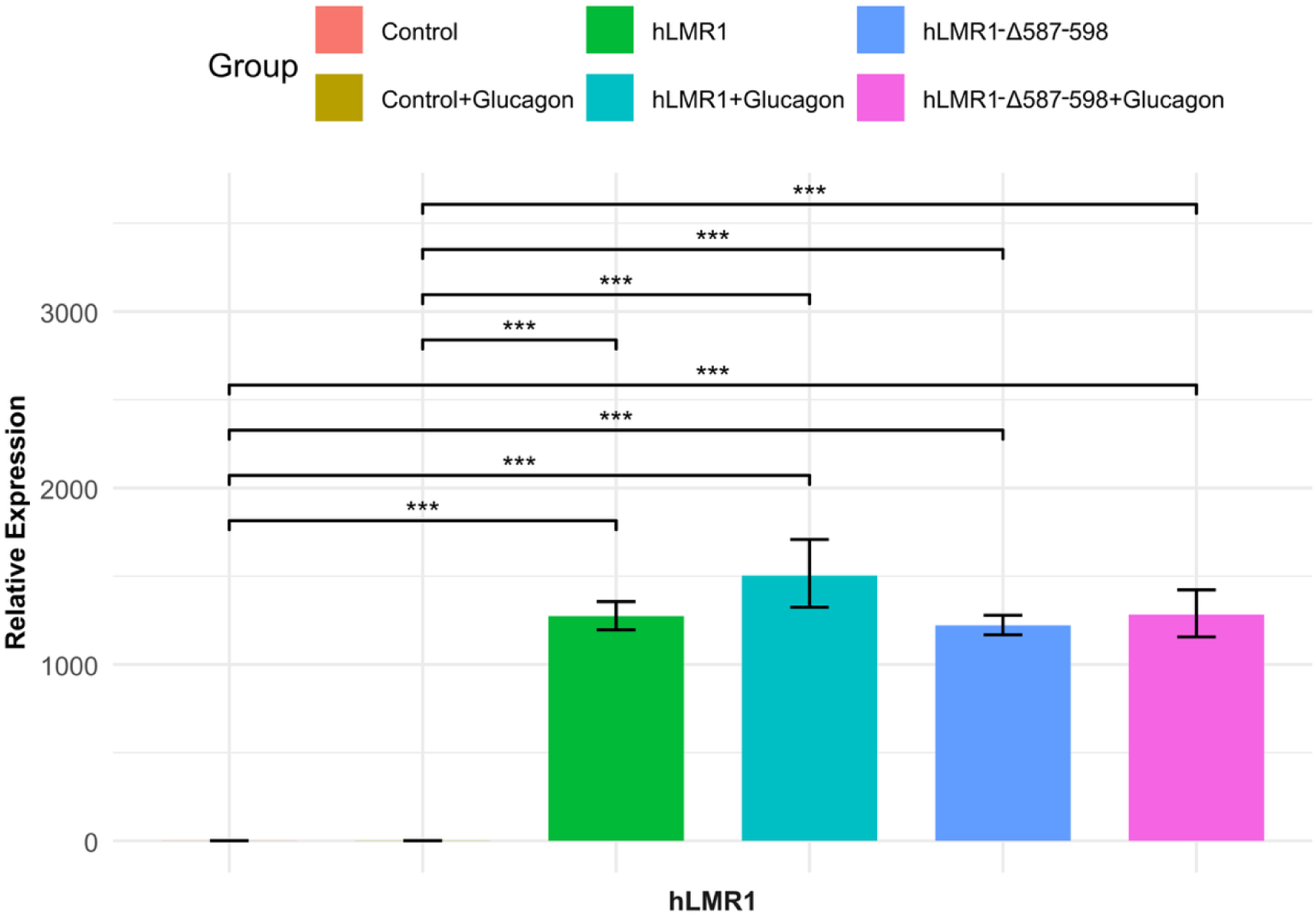
Relative expression of hLMR1 among groups as in Figure 3.

**Supplementary Figure 2.**
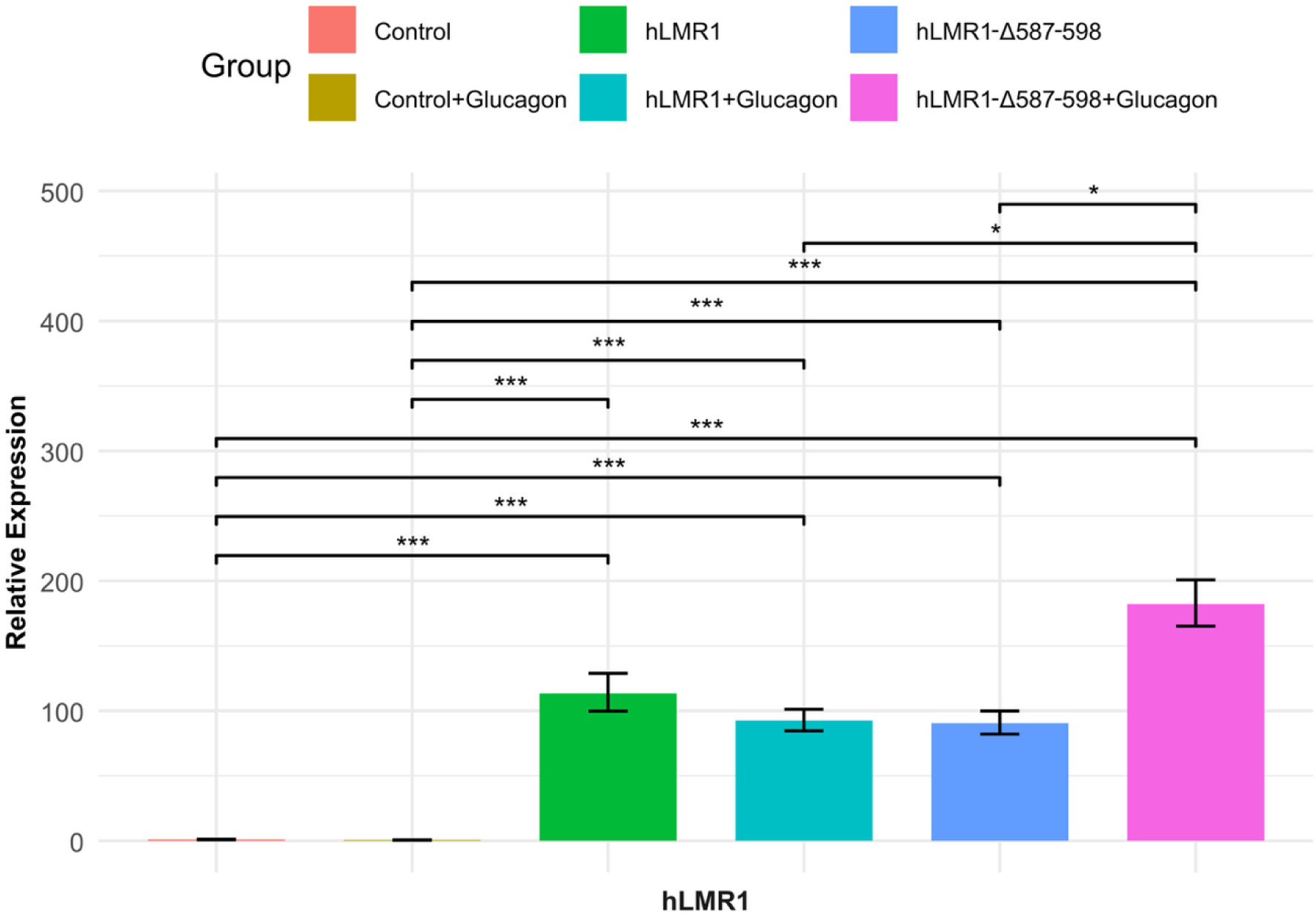
Relative expression of hLMR1 among groups as in Figure 5.

**Supplementary Figure 3.**
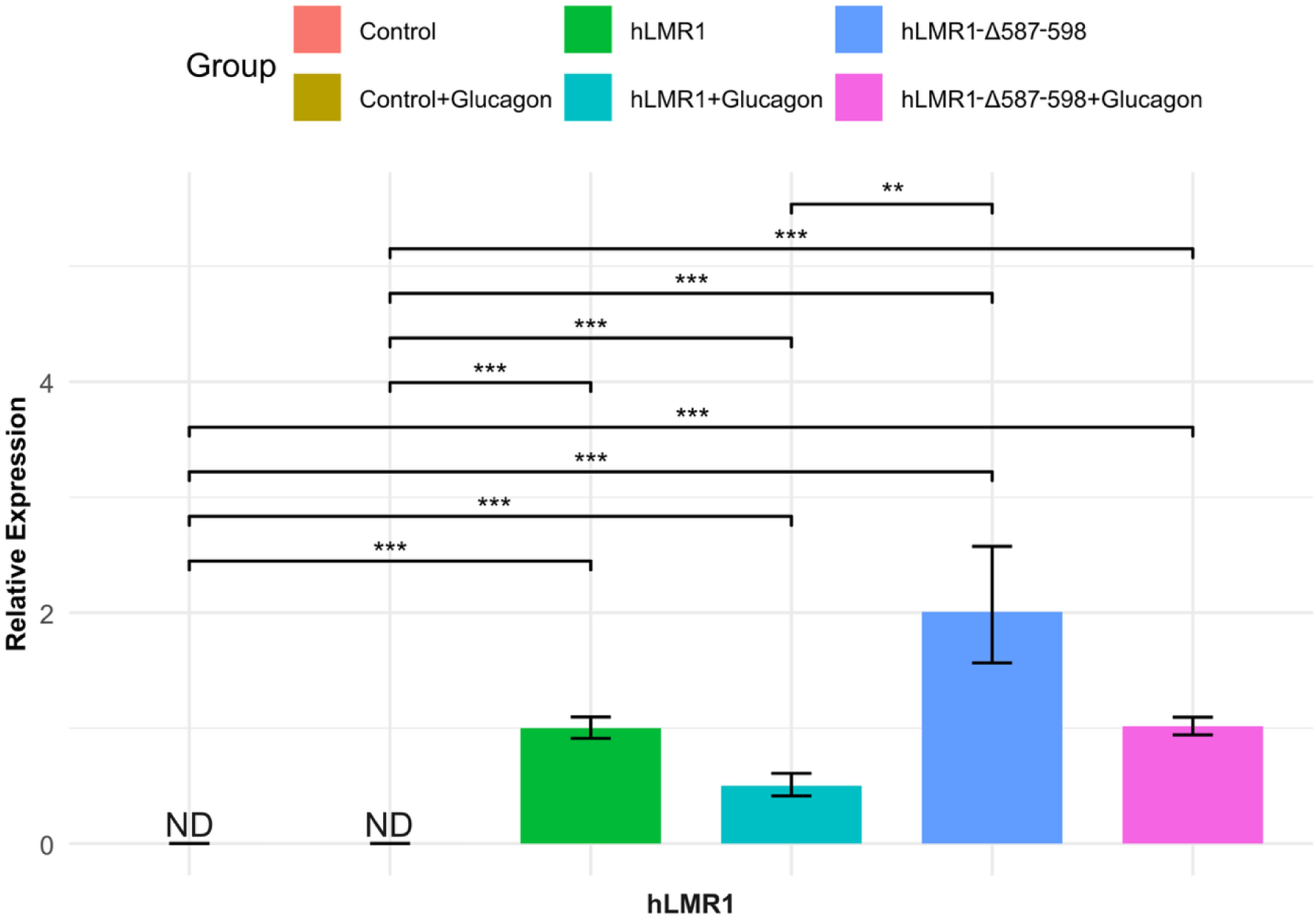
Relative expression of hLMR1 among groups as in Figure 6.

## Table Legend

**Supplementary Table 1.** List of genes that were *differentially regulated* by knocking down of hLMR1 in humanized livers.

**Supplementary Table 2.** List of top 1000 peaks and their associated genes in hLMR1 ChIRP RNA-seq analysis.

**Supplementary Table 3.** Sequences of ChIRP probes, cloning primers, and real-time PCR primers used in this study.

## Author Contributions

M.J. performed the experiments and analyzed the data. L.A.P.F., Z.H., AS-G isolated disease hepatocytes. S.T. performed the bioinformatics analyses and analyzed the data. M.J., S.T. and X.R. wrote the manuscript. X.R. conceived and supervised the study.

## References

1. Kraft, G.; Coate, K. C.; Winnick, J. J.; Dardevet, D.; Donahue, E. P.; Cherrington, A. D.; Williams, P. E.; Moore, M. C., Glucagon’s effect on liver protein metabolism in vivo. Am J Physiol Endocrinol Metab 2017, 313 (3), E263–E272.

2. Eriksen, P. L.; Vilstrup, H.; Rigbolt, K.; Suppli, M. P.; Sorensen, M.; Heeboll, S.; Veidal, S. S.; Knop, F. K.; Thomsen, K. L., Non-alcoholic fatty liver disease alters expression of genes governing hepatic nitrogen conversion. Liver Int 2019, 39 (11), 2094–2101.

3. Gaggini, M.; Carli, F.; Rosso, C.; Buzzigoli, E.; Marietti, M.; Della Latta, V.; Ciociaro, D.; Abate, M. L.; Gambino, R.; Cassader, M.; Bugianesi, E.; Gastaldelli, A., Altered amino acid concentrations in NAFLD: Impact of obesity and insulin resistance. Hepatology 2018, 67 (1), 145–158.

4. Galsgaard, K. D., The Vicious Circle of Hepatic Glucagon Resistance in Non-Alcoholic Fatty Liver Disease. J Clin Med 2020, 9 (12).

5. Hasegawa, T.; Iino, C.; Endo, T.; Mikami, K.; Kimura, M.; Sawada, N.; Nakaji, S.; Fukuda, S., Changed Amino Acids in NAFLD and Liver Fibrosis: A Large Cross-Sectional Study without Influence of Insulin Resistance. Nutrients 2020, 12 (5).

6. Pyo, J. J.; Choi, Y., Key hepatic signatures of human and mouse nonalcoholic steatohepatitis: A transcriptome-proteome data meta-analysis. Front Endocrinol (Lausanne) 2022, 13, 934847.

7. Dean, E. D.; Li, M.; Prasad, N.; Wisniewski, S. N.; Von Deylen, A.; Spaeth, J.; Maddison, L.; Botros, A.; Sedgeman, L. R.; Bozadjieva, N.; Ilkayeva, O.; Coldren, A.; Poffenberger, G.; Shostak, A.; Semich, M. C.; Aamodt, K. I.; Phillips, N.; Yan, H.; Bernal-Mizrachi, E.; Corbin, J. D.; Vickers, K. C.; Levy, S. E.; Dai, C.; Newgard, C.; Gu, W.; Stein, R.; Chen, W.; Powers, A. C., Interrupted Glucagon Signaling Reveals Hepatic alpha Cell Axis and Role for L-Glutamine in alpha Cell Proliferation. Cell Metab 2017, 25 (6), 1362–1373 e5.

8. Solloway, M. J.; Madjidi, A.; Gu, C.; Eastham-Anderson, J.; Clarke, H. J.; Kljavin, N.; Zavala-Solorio, J.; Kates, L.; Friedman, B.; Brauer, M.; Wang, J.; Fiehn, O.; Kolumam, G.; Stern, H.; Lowe, J. B.; Peterson, A. S.; Allan, B. B., Glucagon Couples Hepatic Amino Acid Catabolism to mTOR-Dependent Regulation of alpha-Cell Mass. Cell Rep 2015, 12 (3), 495–510.

9. Suppli, M. P.; Bagger, J. I.; Lund, A.; Demant, M.; van Hall, G.; Strandberg, C.; Konig, M. J.; Rigbolt, K.; Langhoff, J. L.; Wewer Albrechtsen, N. J.; Holst, J. J.; Vilsboll, T.; Knop, F. K., Glucagon Resistance at the Level of Amino Acid Turnover in Obese Subjects With Hepatic Steatosis. Diabetes 2020, 69 (6), 1090–1099.

10. Hennessy, E. J.; van Solingen, C.; Scacalossi, K. R.; Ouimet, M.; Afonso, M. S.; Prins, J.; Koelwyn, G. J.; Sharma, M.; Ramkhelawon, B.; Carpenter, S.; Busch, A.; Chernogubova, E.; Matic, L. P.; Hedin, U.; Maegdefessel, L.; Caffrey, B. E.; Hussein, M. A.; Ricci, E. P.; Temel, R. E.; Garabedian, M. J.; Berger, J. S.; Vickers, K. C.; Kanke, M.; Sethupathy, P.; Teupser, D.; Holdt, L. M.; Moore, K. J., The long noncoding RNA CHROME regulates cholesterol homeostasis in primate. Nat Metab 2019, 1 (1), 98–110.

11. Li, P.; Ruan, X.; Yang, L.; Kiesewetter, K.; Zhao, Y.; Luo, H.; Chen, Y.; Gucek, M.; Zhu, J.; Cao, H., A liver-enriched long non-coding RNA, lncLSTR, regulates systemic lipid metabolism in mice. Cell Metab 2015, 21 (3), 455–67.

12. Mi, L.; Zhao, X. Y.; Li, S.; Yang, G.; Lin, J. D., Conserved function of the long noncoding RNA Blnc1 in brown adipocyte differentiation. Mol Metab 2017, 6 (1), 101–110.

13. Ruan, X.; Li, P.; Cangelosi, A.; Yang, L.; Cao, H., A Long Non-coding RNA, lncLGR, Regulates Hepatic Glucokinase Expression and Glycogen Storage during Fasting. Cell Rep 2016, 14 (8), 1867–75.

14. Ruan, X.; Li, P.; Chen, Y.; Shi, Y.; Pirooznia, M.; Seifuddin, F.; Suemizu, H.; Ohnishi, Y.; Yoneda, N.; Nishiwaki, M.; Shepherdson, J.; Suresh, A.; Singh, K.; Ma, Y.; Jiang, C. F.; Cao, H., In vivo functional analysis of non-conserved human lncRNAs associated with cardiometabolic traits. Nat Commun 2020, 11 (1), 45.

15. Ruan, X.; Li, P.; Ma, Y.; Jiang, C. F.; Chen, Y.; Shi, Y.; Gupta, N.; Seifuddin, F.; Pirooznia, M.; Ohnishi, Y.; Yoneda, N.; Nishiwaki, M.; Dumbovic, G.; Rinn, J. L.; Higuchi, Y.; Kawai, K.; Suemizu, H.; Cao, H., Identification of human long noncoding RNAs associated with nonalcoholic fatty liver disease and metabolic homeostasis. J Clin Invest 2021, 131 (1).

16. Sallam, T.; Jones, M.; Thomas, B. J.; Wu, X.; Gilliland, T.; Qian, K.; Eskin, A.; Casero, D.; Zhang, Z.; Sandhu, J.; Salisbury, D.; Rajbhandari, P.; Civelek, M.; Hong, C.; Ito, A.; Liu, X.; Daniel, B.; Lusis, A. J.; Whitelegge, J.; Nagy, L.; Castrillo, A.; Smale, S.; Tontonoz, P., Transcriptional regulation of macrophage cholesterol efflux and atherogenesis by a long noncoding RNA. Nat Med 2018, 24 (3), 304–312.

17. Schmidt, E.; Dhaouadi, I.; Gaziano, I.; Oliverio, M.; Klemm, P.; Awazawa, M.; Mitterer, G.; Fernandez-Rebollo, E.; Pradas-Juni, M.; Wagner, W.; Hammerschmidt, P.; Loureiro, R.; Kiefer, C.; Hansmeier, N. R.; Khani, S.; Bergami, M.; Heine, M.; Ntini, E.; Frommolt, P.; Zentis, P.; Orom, U. A.; Heeren, J.; Bluher, M.; Bilban, M.; Kornfeld, J. W., LincRNA H19 protects from dietary obesity by constraining expression of monoallelic genes in brown fat. Nat Commun 2018, 9 (1), 3622.

18. Yang, L.; Li, P.; Yang, W.; Ruan, X.; Kiesewetter, K.; Zhu, J.; Cao, H., Integrative Transcriptome Analyses of Metabolic Responses in Mice Define Pivotal LncRNA Metabolic Regulators. Cell Metab 2016, 24 (4), 627–639.

19. Zhao, X. Y.; Li, S.; Wang, G. X.; Yu, Q.; Lin, J. D., A long noncoding RNA transcriptional regulatory circuit drives thermogenic adipocyte differentiation. Mol Cell 2014, 55 (3), 372–82.

20. Johnsson, P.; Lipovich, L.; Grander, D.; Morris, K. V., Evolutionary conservation of long non-coding RNAs; sequence, structure, function. Biochim Biophys Acta 2014, 1840 (3), 1063–71.

21. Hasegawa, M.; Kawai, K.; Mitsui, T.; Taniguchi, K.; Monnai, M.; Wakui, M.; Ito, M.; Suematsu, M.; Peltz, G.; Nakamura, M.; Suemizu, H., The reconstituted ‘humanized liver’ in TK-NOG mice is mature and functional. Biochem Biophys Res Commun 2011, 405 (3), 405–10.

22. Brancale, J.; Vilarinho, S., A single cell gene expression atlas of 28 human livers. J Hepatol 2021, 75 (1), 219–220.

23. Chu, C.; Qu, K.; Zhong, F. L.; Artandi, S. E.; Chang, H. Y., Genomic maps of long noncoding RNA occupancy reveal principles of RNA-chromatin interactions. Mol Cell 2011, 44 (4), 667–78.

24. Gerhard, G. S.; Legendre, C.; Still, C. D.; Chu, X.; Petrick, A.; DiStefano, J. K., Transcriptomic Profiling of Obesity-Related Nonalcoholic Steatohepatitis Reveals a Core Set of Fibrosis-Specific Genes. J Endocr Soc 2018, 2 (7), 710–726.

25. Mardinoglu, A.; Wu, H.; Bjornson, E.; Zhang, C.; Hakkarainen, A.; Rasanen, S. M.; Lee, S.; Mancina, R. M.; Bergentall, M.; Pietilainen, K. H.; Soderlund, S.; Matikainen, N.; Stahlman, M.; Bergh, P. O.; Adiels, M.; Piening, B. D.; Graner, M.; Lundbom, N.; Williams, K. J.; Romeo, S.; Nielsen, J.; Snyder, M.; Uhlen, M.; Bergstrom, G.; Perkins, R.; Marschall, H. U.; Backhed, F.; Taskinen, M. R.; Boren, J., An Integrated Understanding of the Rapid Metabolic Benefits of a Carbohydrate-Restricted Diet on Hepatic Steatosis in Humans. Cell Metab 2018, 27 (3), 559–571 e5.

26. Wewer Albrechtsen, N. J.; Junker, A. E.; Christensen, M.; Haedersdal, S.; Wibrand, F.; Lund, A. M.; Galsgaard, K. D.; Holst, J. J.; Knop, F. K.; Vilsboll, T., Hyperglucagonemia correlates with plasma levels of non-branched-chain amino acids in patients with liver disease independent of type 2 diabetes. Am J Physiol Gastrointest Liver Physiol 2018, 314 (1), G91–G96.

27. Jaso-Vera, M. E.; Takaoka, S.; Patel, I.; Ruan, X., Integrative regulation of hLMR1 by dietary and genetic factors in nonalcoholic fatty liver disease and hyperlipidemia. Hum Genet 2024, 143 (7), 897–906.

28. Lee, B.; Sahoo, A.; Marchica, J.; Holzhauser, E.; Chen, X.; Li, J. L.; Seki, T.; Govindarajan, S. S.; Markey, F. B.; Batish, M.; Lokhande, S. J.; Zhang, S.; Ray, A.; Perera, R. J., The long noncoding RNA SPRIGHTLY acts as an intranuclear organizing hub for pre-mRNA molecules. Sci Adv 2017, 3 (5), e1602505.

29. Kaffe, E.; Roulis, M.; Zhao, J.; Qu, R.; Sefik, E.; Mirza, H.; Zhou, J.; Zheng, Y.; Charkoftaki, G.; Vasiliou, V.; Vatner, D. F.; Mehal, W. Z.; AlcHepNet; Yuval, K.; Flavell, R. A., Humanized mouse liver reveals endothelial control of essential hepatic metabolic functions. Cell 2023, 186 (18), 3793–3809 e26.

30. Ma, J.; Tan, X.; Kwon, Y.; Delgado, E. R.; Zarnegar, A.; DeFrances, M. C.; Duncan, A. W.; Zarnegar, R., A Novel Humanized Model of NASH and Its Treatment With META4, A Potent Agonist of MET. Cell Mol Gastroenterol Hepatol 2022, 13 (2), 565–582.

31. Minniti, M. E.; Foquet, L.; Pedrelli, M.; Gierer, E.; Goldman, D.; Copenhaver, R.; Luquet, S. H.; Grompe, M.; Paolo, P., Liver-Humanized Mice Provide a New Platform to Study Human Cardiometabolic Diseases. Circulation 2021, 144.

32. Bolger, A. M.; Lohse, M.; Usadel, B., Trimmomatic: a flexible trimmer for Illumina sequence data. Bioinformatics 2014, 30 (15), 2114–20.

33. Dobin, A.; Davis, C. A.; Schlesinger, F.; Drenkow, J.; Zaleski, C.; Jha, S.; Batut, P.; Chaisson, M.; Gingeras, T. R., STAR: ultrafast universal RNA-seq aligner. Bioinformatics 2013, 29 (1), 15–21.

34. Danecek, P.; Bonfield, J. K.; Liddle, J.; Marshall, J.; Ohan, V.; Pollard, M. O.; Whitwham, A.; Keane, T.; McCarthy, S. A.; Davies, R. M.; Li, H., Twelve years of SAMtools and BCFtools. Gigascience 2021, 10 (2).

35. Liao, Y.; Smyth, G. K.; Shi, W., The Subread aligner: fast, accurate and scalable read mapping by seed-and-vote. Nucleic Acids Res 2013, 41 (10), e108.

36. Kanehisa, M.; Goto, S., KEGG: kyoto encyclopedia of genes and genomes. Nucleic Acids Res 2000, 28 (1), 27–30.

37. Chu, C.; Quinn, J.; Chang, H. Y., Chromatin isolation by RNA purification (ChIRP). J Vis Exp 2012, (61).

38. Chen, S.; Zhou, Y.; Chen, Y.; Gu, J., fastp: an ultra-fast all-in-one FASTQ preprocessor. Bioinformatics 2018, 34 (17), i884–i890.

39. Zhang, Y.; Liu, T.; Meyer, C. A.; Eeckhoute, J.; Johnson, D. S.; Bernstein, B. E.; Nusbaum, C.; Myers, R. M.; Brown, M.; Li, W.; Liu, X. S., Model-based analysis of ChIP-Seq (MACS). Genome Biol 2008, 9 (9), R137.

40. Quinlan, A. R.; Hall, I. M., BEDTools: a flexible suite of utilities for comparing genomic features. Bioinformatics 2010, 26 (6), 841–2.

41. Heinz, S.; Benner, C.; Spann, N.; Bertolino, E.; Lin, Y. C.; Laslo, P.; Cheng, J. X.; Murre, C.; Singh, H.; Glass, C. K., Simple combinations of lineage-determining transcription factors prime cis-regulatory elements required for macrophage and B cell identities. Mol Cell 2010, 38 (4), 576–89.

42. Ortiz, K.; Cetin, Z.; Sun, Y.; Hu, Z.; Kurihara, T.; Tafaleng, E. N.; Florentino, R. M.; Ostrowska, A.; Soto-Gutierrez, A.; Faccioli, L. A. P., Human Hepatocellular response in Cholestatic Liver Diseases. Organogenesis 2023, 19 (1), 2247576.

